# A restriction enzyme reduced representation sequencing approach for low-cost, high-throughput metagenome profiling

**DOI:** 10.1101/694133

**Authors:** Melanie K. Hess, Suzanne J. Rowe, Tracey C. Van Stijn, Hannah M. Henry, Sharon M. Hickey, Rudiger Brauning, Alan F. McCulloch, Andrew S. Hess, Michelle R. Kirk, Sandra Kittelmann, Graham R. Wood, Peter H. Janssen, John C. McEwan

**Author notes:** Corresponding author (MKH).

## Abstract

Microbial community profiles have been associated with a variety of traits, including methane emissions in livestock, however, these profiles can be difficult and expensive to obtain for thousands of samples. The objective of this work was to develop a low-cost, high-throughput approach to capture the diversity of the rumen microbiome. Restriction enzyme reduced representation sequencing (RE-RRS) using *ApeK*I or *Pst*I, and two bioinformatic pipelines (reference-based and reference-free) were compared to 16S rRNA gene sequencing using repeated samples collected two weeks apart from 118 sheep that were phenotypically extreme (60 high and 58 low) for methane emitted per kg dry matter intake (n=236). DNA was extracted from freeze-dried rumen samples using a phenol chloroform and bead-beating protocol prior to sequencing. The resulting sequences were used to investigate the repeatability of the rumen microbial community profiles, the effect of host genetics, laboratory and analytical method, and the genetic and phenotypic correlations with methane production. The results suggested that the best method was *Pst*I RE-RRS analyzed with the reference-free approach via a correspondence analysis, with estimates for repeatability of 0.62±0.06, heritability 0.31±0.29, and genetic and phenotypic correlation with methane emissions of 0.88±0.25 and 0.64±0.05 respectively for the first component of correspondence analysis. The reference-free approach assigned 62.0±5.7% of reads to common 65 bp tags, much higher than the reference-based approach of 6.8±1.8% of reads assigned. Sensitivity studies suggested approximately 2000 samples could be sequenced in a single lane on an Illumina HiSeq 2500, therefore the current work of 118 samples/lane and future proposed 384 samples/lane are well within that threshold. Our approach is now being used to investigate host factors affecting the rumen and its association with a variety of production and environmental traits. With minor adaptations, our approach could be used to obtain microbial profiles from other metagenomic samples.

## Introduction

Metagenomics is the study of genetic material recovered directly from environmental samples and captures the myriad of organisms present in that environment. Samples of soil and water are obvious examples of environmental samples; however, the gut can also be considered an environment due to the presence of microbes that interact with the host, e.g. during digestion. Metagenomic studies have gained popularity in recent years, primarily in human health, e.g. Irritable Bowel Disease (1) and Coeliac Disease (2).

In an agricultural setting, rumen microbial community (RMC) profiles have been associated with environmentally and economically important traits, such as methane emissions (3, 4) and feed efficiency (5, 6). The RMC breaks down ingested feed to produce volatile or short chain fatty acids, which are a source of energy for the host. Hydrogen produced by this process is metabolized into methane by methanogenic archaea. Methane production is not solely dependent on the abundance of methanogens present, but also on the substrate available to them (7). Different “ruminotypes”, generalized microbial community types, can be found in sheep with high and low methane production, with at least two ruminotypes present in low-methane sheep fed lucerne pellets (3). Furthermore, RMC profiles from rumen samples are moderately heritable (8), suggesting that selection on microbiomes is likely to result in changes in offspring microbiomes. Given that traits such as methane emissions and feed efficiency are difficult and expensive to measure, selection on RMC profiles may facilitate progress in these traits, provided costs are low enough and the method is high-throughput.

Historically, there have been two approaches used for sequencing metagenome samples: targeted sequencing and whole genome shotgun (WGS) sequencing. Targeted sequencing amplifies specified phylogenetically informative genes from a sample, such as the 16S rRNA gene (16S) of microbes, which typically distinguishes taxonomic groups well due to large, comprehensive databases of 16S rRNA sequences that include both culturable and unculturable organisms (9, 10). This approach usually relies on having long sequence reads (11), only captures phylogenetic variation at one gene, and is subject to PCR primer bias due to mismatches in the flanking regions where the primers bind (12). WGS can capture any part of the microbial, host or feed genome; but to be informative, a reference database of genome assemblies with known taxonomies is needed to obtain taxonomic information on WGS sequences, e.g. the Hungate1000 Collection (13). Whole genome assemblies are currently difficult to obtain on unculturable microbes, so these are usually missing from reference databases. Hundreds of millions of reads are generated per sample for WGS, making it an expensive and time-consuming method that additionally requires significant computation resources.

Restriction Enzyme-Reduced Representation Sequencing (RE-RRS, also known as Genotyping-by-Sequencing or GBS) is a next-generation sequencing technique that reduces genome complexity by digestion of genomic DNA by restriction enzymes, followed by the sequencing of fragments within a given size range (14). RE-RRS is used to obtain genotypes for parentage identification or genomic selection (to identify the individuals with the most favorable phenotypes) in a variety of species across livestock, plants and aquaculture (15-18), as well as population diversity studies, e.g. for conservation (19). RE-RRS holds promise as a technique for rapid, high-throughput and cost-effective sequencing of metagenome samples at a fraction of the cost of WGS. Underlying the RE-RRS method is the assumption that sequencing only a specific fraction, typically 0.5-1% of any microbial genome as defined by restriction site and fragment size, captures the majority of information on composition and diversity of the microbial community at a fraction of the sequencing cost. This study used sheep rumen samples to show the potential of RE-RRS as a low-cost, high-throughput approach for obtaining metagenome profiles on thousands of samples, and describes pipelines for obtaining profiles both with and without a reference database.

## Materials and methods

### Rumen sampling and associated methane yields

The sheep rumen samples and methane yield data used for this study were those for which rumen microbial community structure was analysed using 16S rRNA gene sequencing in Kittelmann et al. (3) and part of a larger experiment described in Pinares-Patiño et al. (20). Briefly, respiration chambers were used to measure methane yield (g CH_4_/ kg DMI) on 340 sheep at two independent measuring rounds two weeks apart, each over two days in 4 separate cohorts of animals. The rumen sample was collected via stomach tubing at the end of each measuring round and immediately stored at −20°C. Two rumen samples from a subsample of 118 sheep, representing the ∼17% highest and lowest emitters (60 high-methane and 58 low-methane based on methane yield phenotype), were previously freeze dried, homogenized and stored at −85°C. Subsamples were used for analysis of RMC by sequencing amplified bacterial 16S rRNA genes, and these sequences are available in the EMBL database under the study accession number ERP003779 (3). The 16S rRNA gene profiles used in our study used the recently updated taxonomic classification of these sequences by Kumar (21). These same freeze-dried and homogenised samples were used in our study to evaluate the potential of using RE-RRS for RMC profiling, as described below.

### DNA extraction and Restriction Enzyme-Reduced Representation Sequencing

DNA was extracted from rumen samples using a combined bead-beating, phenol and column purification protocol using the QIAquick 96 PCR purification kit (Qiagen, Hilden, Germany), as described in Text S1 of Kittelmann et al. (3), to provide high quality nucleic acids for RE-RRS. *Ape*KI and *Pst*I restriction enzymes were used separately to test whether RE-RRS is a suitable approach for rumen metagenome profiling. These two enzymes were selected because an in-silico digestion and size filtering of rumen microbial genome assemblies from the Hungate1000 Collection (13) showed that RE-RRS using either *Ape*KI (G|CWGC) or *Pst*I (CTGCA|G) captured microbial sequence from all species present in the collection with an average of 8.6% and 0.3% of each genome, respectively (22).

After digestion of DNA by either *Ape*KI or *Pst*I, barcodes were ligated to link sequences to samples, as described by Elshire et al. (14), and samples were grouped into two libraries, one library for each restriction enzyme used, and PCR amplified. Amplified sequences between 193 and 318 bp (equivalent to 65 – 195 bp inserts) were selected using a Pippin Prep (SAGE Science, Massachusetts, USA) and each library was run on two lanes on the same flow cell on an Illumina HiSeq2500 machine, generating 101 bp single end reads using version 4 chemistry. One plate of 94 samples for *Pst*I were re-run (in a single lane) because barcodes did not ligate in the initial run. FastQ files were deposited in the NCBI SRA.

### Bioinformatic pipeline

Sequenced reads were demultiplexed using GBSX (23), and trimmed using trim_galore for (24) single reads with a minimum length of 40 base pairs. Samples with fewer than 100,000 reads across both lanes of sequencing for a single restriction enzyme were removed from all further analyses, consisting of one sample for *Ape*KI and two samples for *Pst*I. The trimmed sequences from the remaining samples were run through both the reference-based and reference-free pipelines, described below.

#### Reference-based approach

The reference-based (RB) approach used nucleotide BLAST (BLASTN) in BLAST v2.2.28+ (25) with default parameters to compare sequenced reads against the 410 rumen microbial genome assemblies from the Hungate1000 Collection (13). This was found to be the optimal approach for aligning query sequences to the Hungate1000 Collection by Hess et al. (22). Reads were assigned to a taxonomic node using the algorithm from MEGAN (26) implemented in R with default parameters: a minimum bitscore of 50 and considering only hits within 10% of the maximum bitscore for a query read. This approach was evaluated by Hess, Rowe (22) and found to assign reads at the genus level with high accuracy (>96%). The RMC profile was defined as the number of sequences assigned to each of the ∼60 genera represented in the Hungate1000 Collection and associated analyses will be denoted by *Ape*KI_RB and *Pst*I_RB for analyses of *Ape*KI and *Pst*I profiles, respectively.

#### Reference-free approach

The reference-free approach involved collating a set of “tags”, i.e. non-redundant 65 bp-long DNA sequences (evaluated across all samples) commencing at the initial cut site. Tags were required to be present in 10% and 25% of samples in *Ape*KI and *Pst*I, respectively. A comparison of performance for other tag lengths and a variety of prevalence thresholds can be found in File S1– requiring 65 bp tags to be present in at least 10% of samples for *Ape*KI and 25% of samples for *Pst*I gave high estimates of heritability and repeatability with low standard errors, so these parameters were selected for further analysis. The rumen community profile for each sample was generated by counting the abundance of each tag from the sequenced reads; these profiles were collated into a count matrix with samples as rows and tags as columns, obtained using an in-house Unix script. Reference-free analyses will be denoted as *Ape*KI_RF and *Pst*I_RF for profiles from the *Ape*KI and *Pst*I restriction enzymes, respectively.

### Comparison of methods for obtaining RMC profiles

#### Correspondence Analysis

A correspondence analysis (27) was used to reduce the dimensionality of the dataset and facilitate comparisons between the different methods for generating RMC profiles. In a similar approach to that used by Rowe et al. (8), the coordinates of the first dimension of the correspondence analysis (CA1) were used as the RMC profile phenotype for parameter estimation, described below. The first dimension of a correspondence analysis is the dimension that explains the largest proportion of variation in the data.

#### Parameter Estimation

Heritability, repeatability, and genetic and phenotypic correlations with scaled methane yield were estimated in ASReml 4.1 (28). Heritability and repeatability of CA1 were estimated using a univariate mixed linear model, and correlation between CA1 and scaled methane yield was estimated with a bivariate mixed linear model. Scaled methane yield was obtained by dividing methane yield by the contemporary group mean and multiplying by the overall mean, where contemporary group included recording year, lot, group and round and the overall mean was 16.0 kg, as described in Pinares-Patiño et al. (20). In both univariate and bivariate models, sex and cohort (lot and round) were fitted as fixed class effects, random animal genetic effects were estimated based on pedigree relationships, and a random permanent environmental effect linked duplicate samples from the same animal. In some cases, the model was unable to separate animal and permanent environmental effects, in which case animal was dropped from the model and only repeatability reported.

### Sensitivity to sequencing depth

Sequencing more samples per lane would lower the cost of RE-RRS profiling but would consequently reduce the sequencing depth. At low depths the profiling would not accurately capture the proportion of each microbe in the sample, particularly microbes that are in low abundance. Therefore, a sensitivity analysis was performed evaluate the impact of reducing the sequencing depth in our approach. Reads were subsampled with probability 0.5, 0.25, 0.1, 0.05, 0.01, 0.005, 0.002 or 0.001; representing sequencing 2, 4, 10, 20, 100, 200, 500 or 1000 times the number of samples per lane, respectively. The set of sampled reads at a given simulated sequencing depth were then used to calculate repeatability, as above, or compression efficiency (29). Compression efficiency compares the size of a compressed file to its original size as (original – compressed)/original and is a measure of the non-redundant information present in the file. In our study, the original file contained the reads for a given sample without their identifiers. This file was compressed using gzip 1.3.12 (30), which uses the DEFLATE algorithm (31). The value of compression efficiency was the mean across all samples for the simulated sequencing depth. Standard errors for repeatability and compression efficiency were the standard deviation across five replicates at that sequencing depth.

## Results and Discussion

### Sequencing Results

Sequence read quality was high for all lanes sequenced (Figure S1). A greater average number of reads per sample was observed for samples digested with *Pst*I (Table 1), likely partially due to re-running of samples – only 94 samples were run in that lane rather than 118 (i.e. 236 samples across 2 lanes). Sequences from the *Ape*KI digest were slightly shorter than from the *Pst*I digest, however this difference was not significant based on a t-test with alpha = 0.05.

**Table 1:** Average Number of Reads per Sample and Average Read Length of RE-RRS Reads.

| Restriction enzyme | Reads per sample (sd) | Read length (sd)<sup>1</sup> |
| --- | --- | --- |
| *Ape*KI | 2.4M (870k) | 71 (17) |
| *Pst*I | 2.7M (680k) | 84 (15) |

1. Trimmed read length in base pairs after barcode removed

### Reference-based approach

Using the MEGAN algorithm on nucleotide BLAST results, 5.3±1.7% and 6.8±1.8% of reads were assigned at the genus level for *Ape*KI and *Pst*I, respectively. This assignment rate is consistent with querying the Hungate1000 Collection with reads from WGS (22, data from Shi et al. (7)). Comparing against a protein database is one method that could potentially improve the proportion of sequences assigned (hit rate) at the genus level. However, Hess et al. (22) found only a small increase in hit rate at the genus level when BLASTX was used. They also found the time taken to perform the BLAST query and analyze the results was much longer when BLASTX was used compared to BLASTN and therefore was not desirable for a high-throughput pipeline.

A significant difference in hit rate between high- and low-methane animals was found for both *Ape*KI (p = 1.4 × 10^−8^) and *Pst*I (p = 1.4 × 10^−8^; Table 2). This may be attributed to the presence or absence of some species associated with methane yield in the Hungate1000 Collection. For example, Kittelmann et al. (3) identified the genera *Fibrobacter, Kandleria, Olsenella* and *Sharpea* to be in higher prevalence in low-methane yield animals. These genera are all present within the Hungate1000 Collection and have equal or significantly higher abundance in samples from low-methane animals. The Hungate1000 Collection also has poor or no representation of other genera that were found by Kittelmann et al. (3) to be in higher abundance in high-methane yield animals *Coprococcus*. This shows that using a method that is reliant on a reference database is limited by the genomes present within that database.

**Table 2:** Hit Rates by Taxonomic Level from RE-RRS Samples using One of Two Restriction Enzymes.

| Restriction enzyme | Sample<sup>1</sup> | Hit rate by taxonomic level (%) |  |  |  |  |  |  |
| --- | --- | --- | --- | --- | --- | --- | --- | --- |
|  |  | Kingdom | Phylum | Class | Order | Family | Genus | Species |
| *ApeKI* | High | 4.9 | 4.9 | 4.8 | 4.8 | 4.7 | 4.7 | 1.4 |
|  | Low | 6.2 | 6.1 | 6.1 | 6.1 | 5.9 | 5.9 | 2.2 |
| *PstI* | High | 6.5 | 6.4 | 6.4 | 6.4 | 6.3 | 6.3 | 1.7 |
|  | Low | 7.4 | 7.4 | 7.4 | 7.4 | 7.3 | 7.3 | 2.3 |

1. Methane yield classification (high- or low-methane yield) of the sheep the sample came from.

A major gap in microbial genome assemblies is the inability, at least historically, to sequence the unculturable microbes that make up a large proportion of any environment. Technological advances, such as single-cell sequencing (32) and the ability to assemble genomes from metagenomic datasets (33), offer alternative solutions to sequence and assemble microbial genomes and will provide opportunities to improve reference databases. Judicious addition of new microbial genome assemblies as they become available will improve hit rates, however, any additional sequences added to the database will also increase the time to complete the analysis, which may not be desirable for a high-throughput approach if there are time constraints. If additional genomes were to be added to the Hungate1000 Collection (or another reference database), expert curation would be needed to ensure the quality of genome assemblies and balance the resource across taxa to maximise coverage and minimise duplication.

### Reference-free approach

The reference-free approach is not subject to the biases of the species represented in the Hungate1000 Collection. We explored the use of different tag lengths and filtering thresholds and showed that there were different optimal filtering levels when using *Ape*KI (10%) or *Pst*I (25%) restriction enzymes (File S1). When *Ape*KI was used, there were ∼1.2M 65 bp tags that were present in at least 10% of the samples and these tags accounted for 20.0±3.5% of reads (corresponding to 33.6±5.8% of reads at least 65 bp long). When *Pst*I was used, there were ∼500k 65 bp tags present in at least 25% of samples and 53.3±5.9% of reads were accounted for (corresponding to 61.9±5.7% of reads at least 65 bp long). Although these proportions are given at different filtering levels, File S1 shows that *Pst*I captures a greater proportion of reads than *Ape*KI at all filtering levels and tag lengths. These differences can be explained by the proportion of each microbial genome that is expected to be captured using each restriction enzyme: *Ape*KI captures 8.6% of Hungate1000 Collection genomes on average, while *Pst*I captures 0.3% (22). This means that, for a given number of sequences (e.g. one lane of sequencing), the fewer regions captured by *Pst*I reads will be at higher depth, whereas the greater number of regions captured by *Ape*KI will be at lower depth. This is shown by the much larger number of unique tags present when using *Ape*KI compared to *Pst*I (File S1) despite a slightly higher number of reads per sample for *Pst*I (Table 1).

The reference-free pipeline is particularly useful for prediction of a trait that is correlated with a microbial profile because knowledge of which taxonomic group a sequence belongs to is of less importance than its predictive ability, so sequences that don’t align to a reference genome can still be utilized. Subsequently, if taxonomic information is desired, tags can be searched against a relevant database; this process would be computationally inexpensive because there are fewer search terms, i.e. fewer tags (hundreds of thousands) than the full set of reads in the original dataset (tens or hundreds of millions). Given the large number of tags generated using the reference-free approach it may be desirable to cluster these into groups (e.g. through sequence similarity, taxonomic assignment, high positive correlations between tag abundances). Tags that come from the same organism will be highly correlated, however, a high correlation (positive or negative) could also come about due to interactions between the microbes, or by chance.

### Comparison of methods for obtaining rumen microbial profiles

#### Variance components of RMC profiles

The first component of the correspondence analysis (CA1) was analysed as a trait for the four RE-RRS approaches and the 16S rRNA gene taxonomic classifications from Kumar (21) on the same samples. The variance of CA1 from the 16S rRNA gene profile was the highest, capturing 24.3% of the variance in that profile (Table 3). The reference-based approaches had the smallest variances, but they explained 40-50% of the variation in the profiles. Lastly, the reference-free approaches both had variances just over 0.2 and explained less than 5% of the variance in these profiles. The percent variance explained by CA1 was negatively correlated with the number of tags or taxa: the reference-based approaches assigned reads to only 60 genera; the 16S rRNA gene approach assigned reads to ∼250 genera and the reference-free approach assigned reads to ∼1.2M and ∼500k tags for *Ape*KI and *Pst*I, respectively. This indicates that a large proportion of the variation in the rumen community profile is not accounted for by analysing only CA1 for these profiles, particularly for the reference-free approach. Nevertheless, evaluating CA1 allowed us to easily compare the different approaches and below we have discussed statistical approaches that will utilize more of the information contained within the profile.

**Table 3:** Variance, heritability and repeatability of the first component of a correspondence analysis of rumen community profiles.

| Method | %Var | Var | Gen Var | Heritability | Repeatability |
| --- | --- | --- | --- | --- | --- |
| 16S | 24.3 | 0.30 | $0.06 \pm 0.05$ | $0.26 \pm 0.23$ | $0.45 \pm 0.08$ |
| *ApeKI*_RB | 47.4 | 0.07 | $0.03 \pm 0.02$ | $0.58 \pm 0.32$ | $0.61 \pm 0.06$ |
| *PstI*_RB | 41.0 | 0.06 | NE | NE | $0.60 \pm 0.06$ |
| *ApeKI*_RF | 3.3 | 0.22 | $0.03 \pm 0.03$ | $0.18 \pm 0.25$ | $0.60 \pm 0.06$ |
| *PstI*_RF | 3.9 | 0.22 | $0.03 \pm 0.04$ | $0.24 \pm 0.27$ | $0.62 \pm 0.06$ |

%Var = Percent of the overall variance explained by first component of correspondence analysis
Var = Variance of first component of correspondence analysis
Gen Var = Genetic variance of first component of correspondence analysis
16S = 16S rRNA gene sequencing approach
RB = Reference-based approach
RF = Reference-free approach
NE = Not Estimable

Despite the differences in the variances of the first component of the correspondence analysis, the genetic variance was very similar for each of the methods (Table 3). The combination of similar genetic variances but different phenotypic variances produced heritability estimates ranging from 0.18 (*Ape*KI_RF) to 0.58 (*Ape*KI_RB), with the 16S rRNA gene approach giving an intermediate heritability estimate of 0.26. Despite coming from animals with extreme phenotypes for methane yield, these heritability estimates are consistent with other studies on livestock and human intestinal microbial communities (4, 8); however, all estimates of heritability in our study had large standard errors due to the relatively small number of samples and were not significantly different from each other. The *Pst*I_RB approach was unable to separate genetic and permanent environmental effects, with the animal effect being bound at zero, so the animal effect was removed from the model. The inability of the *Pst*I_RB approach to separate these effects, as well as the low variance of that component, indicates that it may be a less powerful approach. Increasing the number of samples would allow the model to more accurately separate the genetic and permanent environmental effects for all methods.

Repeatability is a measure of the similarity of two samples from the same individual and is calculated as the genetic plus permanent environmental effect as a proportion of the phenotypic variance. CA1 of the RMC profiles from RE-RRS were more consistent across time than 16S rRNA gene profiles, as evidenced by repeatability estimates almost 1.5 times higher (Table 3).

#### Correlations with methane yield

The genetic correlation of the first component of the correspondence analysis with methane yield was highest for the reference-free approach using *Pst*I, followed by the 16S rRNA gene approach and the reference-based approach using *Ape*KI (Table 4). Estimates were not available for the other two approaches due to an inability to separate genetic and permanent environmental effects. Phenotypic correlations between the first component of a correspondence analysis and methane yield were highest for the two reference-free approaches, followed by the two reference-based and finally the 16S rRNA gene results. These phenotypic and genetic correlations suggest that RE-RRS may produce rumen community profiles with a greater predictive ability than profiles produced from 16S rRNA gene sequencing; however, more samples are needed to test this accurately. The *Ape*KI_RB method has the same genetic correlation with methane yield as 16S rRNA gene sequencing but a higher phenotypic correlation, suggesting that *Ape*KI captures more of the metagenomic variation that is phenotypically correlated with methane yield. There is likely to be some non-microbial DNA that is being captured and influencing the correlations. These results suggest that *Ape*KI will predict the individual’s methane production better than 16S rRNA gene sequencing but not necessarily the genetic potential of that individual as a parent.

**Table 4:** Genetic and phenotypic correlations of the first component of a correspondence analysis of rumen community profiles with methane yield.

| Method | $r_g$ (CH<sub>4</sub> Yield) | $r_p$ (CH<sub>4</sub> Yield) |
| --- | --- | --- |
| 16S | $0.63 \pm 0.49$ | $0.26 \pm 0.07$ |
| *ApeKI*_RB | $0.63 \pm 0.31$ | $0.48 \pm 0.06$ |
| *PstI*_RB | NE | $0.33 \pm 0.07$ |
| *ApeKI*_RF | NE | $0.66 \pm 0.04$ |
| *PstI*_RF | $0.88 \pm 0.25$ | $0.64 \pm 0.05$ |

16S = 16S rRNA gene sequencing approach
RB = Reference-based approach
RF = Reference-free approach
NE = Not Estimable

The higher correlations between methane yield and the reference free approaches (Table 4) suggest that this approach might be capturing components of the rumen microbiome that are not being captured by 16S rRNA gene sequencing or the reference-based approaches. Both 16S rRNA gene and the reference-based approaches are focused on capturing microbial community profiles, while the reference-free approach may capture DNA from a much wider taxonomic range, e.g. host, feed, viruses. If the aim is to obtain the most accurate predictions, then this information from a wider taxonomic range is beneficial to include in the analysis. The reference-free approaches also have the lowest proportion of the variance captured within the first component of the correspondence analysis (Table 3), suggesting that there is potential for the reference-free rumen community profiles to more accurately predict methane yield if additional information is considered. The additional information that is not contained within the first component will be a mixture of “signal” that is associated with methane yield, and “noise” that is not. Too much noise will have a negative impact on prediction accuracies; approaches to separate signal from noise are suggested in the statistical modelling section below.

### Sensitivity to sequencing depth

The number of samples per lane influences the cost per sample of sequencing as well as the average number of sequenced reads per sample. A sensitivity analysis was performed by subsampling reads from our RE-RRS samples and evaluating the repeatability of the first component of a reference-based correspondence analysis, as well as the compression efficiency of the dataset. Both analyses showed that sampling 5% of reads, corresponding to 20 times the number of samples per lane, i.e. 2000 samples, is the lower bound because compression efficiency drops, and the standard error of repeatability increases when sequencing depth is lowered beyond this point (Figure 1).

**Figure 1:**
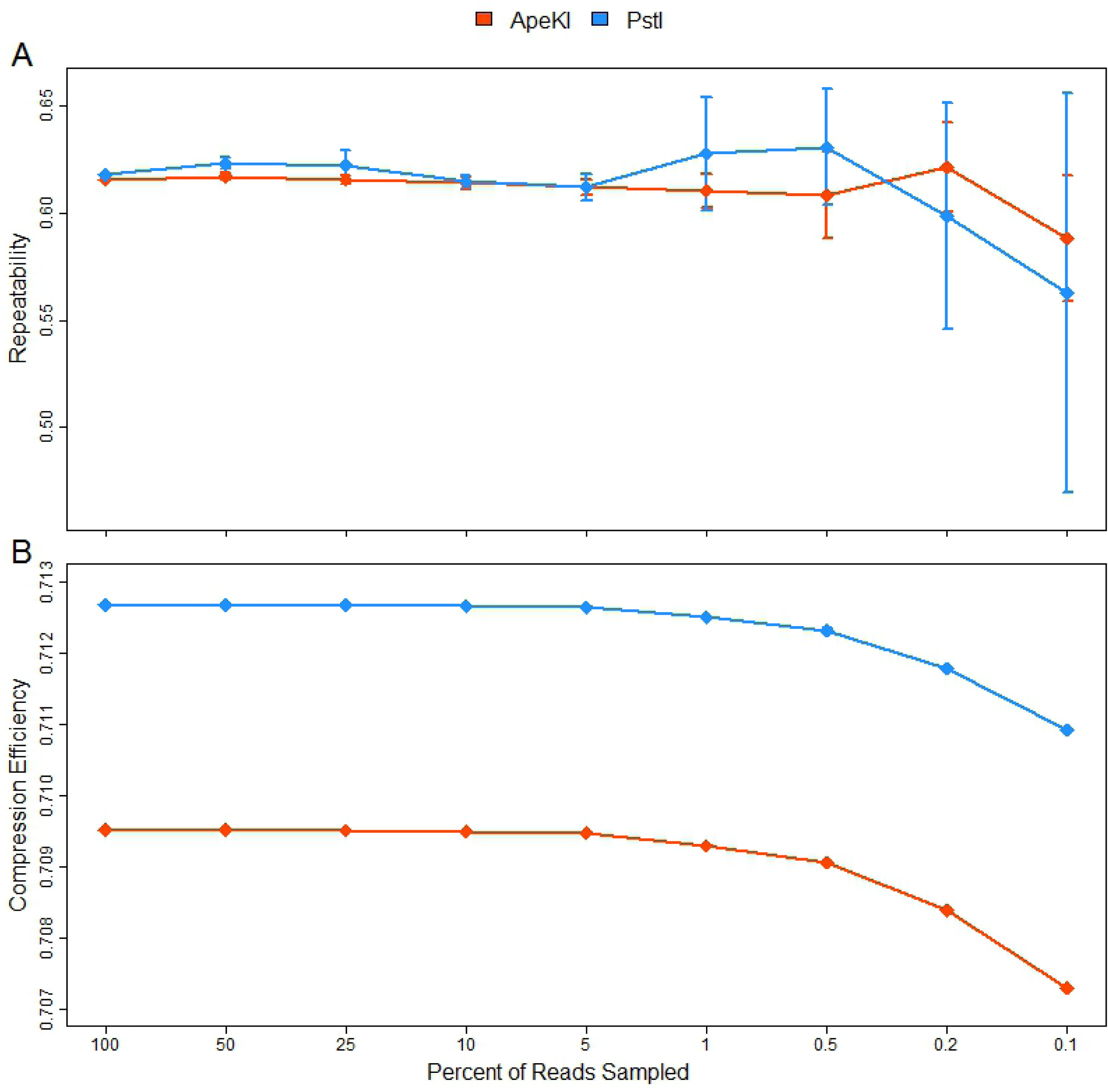
Repeatability (A) and Compression Efficiency (B) of RE-RRS Data as the Percent of Reads Sampled Decreases. The standard error of the estimate of repeatability of the first component of a reference-based correspondence analysis increases (A), and the compression efficiency of sequence data decreases (B), when less than 5% of reads are sampled, a sequencing depth that corresponds to 20 times the number of samples sequenced per lane. This number is consistent across both restriction enzymes used for this study. Standard errors for compression efficiency are negligible and therefore not visible (B).

We have determined a potential cut-off for how much we can increase the throughput without losing crucial information using two separate approaches (compression efficiency and repeatability, Figure 1). It is worth mentioning, however, that these samples are from a relatively small set of individuals that have extreme phenotypes, so in practice more sequences may be needed than the observed cut-off. This analysis shows that the depth with which sequencing has occurred in this study is well within reasonable bounds for capturing metagenomics data, and that the throughput could be safely increased 2-4× over what was done in this study. This will reduce costs and allow faster turn-around times for obtaining sequencing data when large numbers of samples are analysed.

### Utility of a high-throughput Metagenomics Method

#### RE-RRS vs. 16S rRNA gene sequencing and WGS

Our RE-RRS approach to sequencing rumen samples is likely to perform as well or better than 16S rRNA gene sequencing in terms of the variation in sequence reads that is accounted for, and the predictive ability of rumen community profiles (Table 2). Although RE-RRS can capture taxonomic information, like 16S rRNA gene sequencing it cannot directly quantify the abundance of particular genes within a sample (16S rRNA gene sequencing only captures the relative abundance of variants of the 16S rRNA gene); most will be missed because it is a reduced representation sequencing approach that only captures a small percentage of each microbial genome. WGS can capture information on the abundances of these genes; however, sequencing must be done at high depth to capture this information accurately, which is very expensive.

With any metagenomics sequencing approach it is important to try to avoid biases in what is captured. The 16S rRNA gene is present in all bacteria and archaea and contains some highly conserved regions, suitable for universal primer design but also contains highly variable regions which allows the sequences to be assigned to taxonomies. However, sometimes the highly conserved regions have some variation, which leads to primer bias (e.g. Sim et al. (12)). RE-RRS was shown to capture part of each genome in the Hungate1000 Collection (22), suggesting it is not prone to similar biases; however, the Hungate1000 Collection captures only culturable microbes so it is possible that some other microbes are not captured by our approach. This can be evaluated further as more microbial genome assemblies become available.

16S rRNA gene sequencing, WGS and the reference-based RE-RRS approach all require a reference database to assign taxonomic information to the sequences. 16S rRNA gene reference databases tend to be more comprehensive because only a single gene needs to be sequenced, enabling both culturable and unculturable microbes to be captured (10). WGS and the reference-based RE-RRS approach both need reference databases containing genome assemblies. These databases are (currently) less complete and capture a smaller range of taxa. All reference databases such as this are prone to sequencing errors present in the reference database, which may result in incorrect taxonomic assignment.

#### Reference-based vs reference-free

The reference-free approach was developed to overcome the reliance on a reference-database to generate the RMC profile. If the goal of the analysis is prediction of a trait or disease status, it is not critical to know the taxonomic origin of a sequence, as long as it can positively influence prediction accuracy. Our results in Tables 3 and 4 indicate that the reference-free approach will be a valuable approach for predicting methane yield. The reference-free approach is likely capturing more than just microbial variation e.g. host, feed and microbial eukaryotes, which may all play a role in methane emissions.

#### Impact of different restriction enzymes

The choice of restriction enzyme will impact sample throughput. *Ape*KI has a lower compression efficiency than *Pst*I, which is consistent with *Ape*KI capturing more of the genome (Figure 1). This means that two sequences from the same genome with *Pst*I are more likely to be identical (i.e. less of the genome but at a higher depth), resulting in improved compression efficiency. This can also be inferred from the filtering threshold required to achieve a similar repeatability estimate for *Ape*KI (tags must be present in at least 10% of samples, ∼1.2M tags) compared to *Pst*I (tags must be present in 25% of samples, ∼500k tags) (File S1).

The *Pst*I_RF approach performed the best of all the models in terms of heritability, repeatability, and correlations with methane yield. File S1 shows that heritability and repeatability estimates changed very little when different filtering parameters and tag lengths were applied to the *Pst*I_RF data. This indicates that the depth of sequencing is appropriate, which is supported by the higher compression efficiency of *Pst*I compared to *Ape*KI (Fig 1b).

*Ape*KI had a lower compression efficiency than *Pst*I (Fig 1b), and when the reference-free approach was used tags needed to be present in 10% of samples before heritability and repeatability estimates were in line with *Pst*I (File S1). This is because *Ape*KI captures more regions of the microbial genome at lower depth than *Pst*I so is not as powerful for the reference-free approach. By extension, the reference-free approach developed here would not be suitable for WGS data because any part of the metagenome could be captured, this could explain the poor results using a k-mer approach in Ross et al. (34). Therefore, the profiling pipeline needs to be chosen based on the sequencing approach used and the intended analysis.

Reducing tag length will join tags with similar sequences (e.g. two 65 bp tags that differ at only base 40 will be merged into a single 32 bp tag). The shorter tag might represent a higher taxonomic order than the larger one (possibly being less biologically informative, given the higher specificity of a longer sequence) but this is outweighed by the increased power to accurately capture the abundance of that taxonomic group.

These results show the importance of selecting an appropriate restriction enzyme for the purpose of the study when using RE-RRS for metagenome profiling.

#### Application of RE-RRS in livestock

Most metagenome studies in livestock have used small sample sizes and many used animals with extreme phenotypes, which is valuable for identifying whether there is a relationship between microbes and traits of interest. However, knowledge of the RMC of thousands of animals has the potential to reduce the carbon footprint of farming through selection of individuals with a rumen microbiome genetically associated with lower methane emissions. Traits aimed at reducing the carbon footprint of livestock animals, e.g. methane emissions or feed efficiency, are often difficult and expensive to measure. Therefore, provided the costs can be reduced sufficiently and high-throughput profiling is possible, large volumes of samples can be processed and the data analyzed quickly and cheaply. In this situation, metagenome profiling could provide an alternative solution to reducing the carbon footprint that circumvents the need to continually measure expensive methane yield phenotypes on thousands of animals.

#### Other sample types

Much research has been done into sequencing microbial samples from humans (35, 36), particularly samples related to the digestive tract and their association with a variety of health issues (1, 2). A cheap, high-throughput metagenome sequencing approach has the potential to make screening of these samples more accessible to those that require them. High-throughput metagenome profiling has the potential to improve monitoring of other environmental samples as well. This could range from identifying pathogens in water samples, evaluating the quality of water in different environments, to identifying favorable and unfavorable soil environments for the growth of particular crops. Further research is required to evaluate the potential of RE-RRS in each of these situations.

#### Statistical modelling

Biological data has increased in complexity in recent years, as technological advances have enabled the collection of many different parameters on large numbers of samples. Metagenomics provides a rich source of information to include in predictions of associated traits, for example rumen metagenomics samples to predict methane emissions. Statistical methodologies have advanced along with data complexity, enabling the integration of many different –omics in an attempt to improve prediction accuracies. However, it is important to appropriately model this information to obtain robust predictions that extend to a wide diversity of situations.

One important aspect is to appropriately represent the data. Some taxa are present in all samples while others appear in only some samples. For those present in most or all samples, a correlation between the abundance of each tag/genus within each sample is the most appropriate measure; however, for those present in only a subset of samples, a presence/absence coding may be more appropriate. The abundance of each genera within a sample can vary by orders of magnitude, therefore for prediction purposes it is common to transform the counts or proportions of each taxonomic group, often a log transformation (37).

We summarized the RMC profile using the first component of a correspondence analysis; however, other approaches can also be used, e.g. heritability and repeatability could be calculated for each genus or tag. This information can be used to remove genera/tags, or a “metagenome wide association study” (38) – similar to a genome wide association study – can be used to identify tags that are associated with the trait of interest for inclusion in prediction models. This approach will reduce the amount of noise in the profile, hopefully improving prediction accuracy. Other ways to reduce the dimensionality, particularly for the reference-free approach, would be to calculate correlations between each tag and merge the counts for those with high correlations. This approach may be more powerful than using all tags independently because if a tag is not sequenced by chance even though it is present in the sample then a correlated tag may be picked up instead; this will be most important for genera at low abundance.

The metagenomics information can be included in prediction models in a variety of ways depending on the circumstances. Microbial Relationship Matrices (MRMs) have been used to include microbial information to predict traits of interest, particularly in livestock (37). This approach will work when there are more taxa/tags than samples but runs into problems when the opposite is true (39). When there are more samples than taxa/tags, fitting each taxa/tag as a covariate, such as in Bayesian random regression model, is an alternative approach (39). Machine learning algorithms may be useful for integrating host genetic information with microbial data for trait predictions, while random forests hold potential for classifying samples into groups. Our development of a high-throughput, low-cost approach to sequence rumen metagenomics samples has set the stage for generating a large dataset where different modelling approaches can be compared in the future.

## Conclusions

We have shown that RE-RRS is a promising method for obtaining low-cost, high-throughput metagenomic profiles that performs at least as well as 16S rRNA gene sequencing. Metagenomic profiles can be generated either with or without a reference database (reference-based or reference-free, respectively) depending on the purpose of the analysis. Gathering metagenomic information on a large number of animals can be a useful addition to genomic information for the prediction of traits in livestock production and human health. The next steps are to use this approach to sequence thousands of environmental samples and develop appropriate statistical models for prediction purposes.

## Acknowledgements

Our thanks to Drs Graeme Attwood, Kathryn McRae and Ken Dodds (all from AgResearch) for critically reviewing this manuscript, and to Dr Graeme Attwood for insightful discussions throughout the process.

---

**Figure S1: Sequence Quality per Base Pair for all Lanes of Sequencing**

Box and whisker plots of sequence quality (Phred Score) at positions along the sequenced read. Red, orange and g een signify low, medium and high quality bases, respectively. Sequence quality was high throughout the entire read; however, it did drop slightly towards the end of the read. Sequence quality for ApeKI was more variable than for PstI. The third plot for PstI represents the 94 samples that were re-sequenced due to barcodes not ligating in the initial run.

**File S1: Tag Filtering Comparison**

This file contains an investigation into suitable tag lengths (16, 32 or 65 bp) and tag prevalence thresholds (present in 10, 25, 50 or 100% of samples) for use with *Ape*KI and *Pst*I restriction enzymes.

